# Genome-wide association analysis of hyperspectral reflectance data to dissect growth-related traits genetic architecture in maize under inoculation with plant growth-promoting bacteria

**DOI:** 10.1101/2022.08.11.503682

**Authors:** Rafael Massahiro Yassue, Giovanni Galli, Chun-Peng James Chen, Roberto Fritsche-Neto, Gota Morota

**Affiliations:** Department of Genetics, ‘Luiz de Queiroz’ College of Agriculture, University of São Paulo, São Paulo, Brazil; School of Animal Sciences, Virginia Polytechnic Institute and State University, Blacksburg, USA; Center for Advanced Innovation in Agriculture, Virginia Polytechnic Institute and State University, Blacksburg, VA 24061 USA; Quantitative Genetics and Biometrics Cluster, International Rice Research Institute, Los Baños, Philippines

**Keywords:** genome-wide association analysis, growth trait, hyperspectral wavelength, multi-phenotype

## Abstract

Plant growth-promoting bacteria (PGPB) may be of use for increasing crop yield and plant resilience to biotic and abiotic stressors. Using hyperspectral reflectance data to assess growth-related traits may shed light on the underlying genetics as such data can help assess biochemical and physiological traits. This study aimed to integrate hyperspectral reflectance data with genome-wide association analyses to examine maize growth-related traits under PGPB inoculation. A total of 360 inbred maize lines with 13,826 single nucleotide polymorphisms (SNPs) were evaluated with and without PGPB inoculation; 150 hyperspectral wavelength reflectances at 386–1,021 nm and 131 hyperspectral indices were used in the analysis. Plant height, stalk diameter, and shoot dry mass were measured manually. Overall, hyperspectral signatures produced similar or higher genomic heritability estimates than those of manually measured phenotypes, and they were genetically correlated with manually measured phenotypes. Furthermore, several hyperspectral reflectance values and spectral indices were identified by genome-wide association analysis as potential markers for growth-related traits under PGPB inoculation. Eight SNPs were detected, which were associated with manually measured and hyperspectral phenotypes. Moreover, the hyperspectral phenotypes were associated with genes previously reported as candidates for nitrogen uptake efficiency, tolerance to abiotic stressors, and kernel size. In addition, a Shiny web application was developed to explore multi-phenotype genome-wide association results interactively. Taken together, our results demonstrate the usefulness of hyperspectral-based phenotyping for studying maize growth-related traits in response to PGPB inoculation.

## Introduction

Sustainably increasing food production is required considering growing demands, especially in developing nations (Laurance et al., 2014). Recent studies found that plant growthpromoting bacteria (PGPB) are a viable option for increasing plant resilience to biotic and abiotic stressors, with the potential to increase food production (Compant et al., 2005; Batista et al., 2018; Yassue et al., 2021). Plant growth-promoting bacteria can promote morphological (Mantelin, 2003) and functional (Di Benedetto et al., 2017) changes in plants. Reported effects include increased uptakes of nutrients, such as nitrogen, phosphate, potassium, and iron (Egamberdiyeva, 2007; Pii et al., 2015), and activation of responses to pathogens and abiotic stressors (Olanrewaju et al., 2017; Singh et al., 2018).

One of the challenges associated with assessing the possible benefits of PGPB is identifying plant response. Hyperspectral image data can be used to assess biochemical or physiological attributes of plants, thus such data have been increasingly applied in plant genetics and management studies because of their associations with target phenotypes, such as water content (550–1750 nm) (Ge et al., 2016), plant nutrient status (350–2500 nm) (Mahajan et al., 2014; Nigon et al., 2021), disease susceptibility (Thomas et al., 2017), yield (400-900 nm) (Yang et al., 2021), and biomass (380–850 nm) (Krause et al., 2019). For example, hyperspectral patterns genetically correlated with target phenotypes can potentially aid genomic prediction (Krause et al., 2019; Sandhu et al., 2021). In addition, hyperspectral phenotypes can be used for genetic inference studies, such as genome-wide association (GWA), heritability, and genetic correlation analyses, to investigate associations between hyperspectral bands and genome (Feng et al., 2017; Sun et al., 2019; Barnaby et al., 2020; Wu et al., 2021).

Bayesian whole-genome regression models are useful for GWA studies because they implicitly account for population structure and the multiple-testing problem of classical singlemarker linear mixed models by simultaneously fitting all markers (Fernando et al., 2017; Wolc and Dekkers, 2022). Despite the increasing use of high-throughput phenotyping through the compilation of hundreds or thousands of phenotypes, a limited number of whole-genome regression studies have integrated hyperspectral data into genetic inference research. Moreover, how hyperspectral wavelength data are associated with PGPB responses in maize remains elusive because it is challenging to interpret changes in hyperspectral reflectance patterns with regard to plant biological processes. The objectives of this study were 1) to investigate whether hyperspectral reflectance values are under genomic control, 2) to determine whether variations in hyperspectral reflectance values are correlated with growth-related traits at the genomic level, and 3) to identify specific genomic regions associated with wavelengths that can be used to study maize growth-related traits under PGPB inoculation. We employed Bayesian whole-genome GWA methods to identify possible candidate genes associated with growth-related traits and hyperspectral reflectance bands. In addition, a Shiny web application was developed to to explore the multi-phenotype GWA results interactively.

## Materials and Methods

### Plant growth-promoting bacteria experiment

A tropical maize association panel comprising 360 inbred lines was used to examine genetic basis of PGPB responses. The inbred lines were evaluated under nitrogen stress with (B+) and without (B-) PGPB inoculation. Maize seeds were co-inoculated with the PGPB strains *Bacillus thuringiensis* RZ2MS9, *Delftia* sp. RZ4MS18 (Batista et al., 2018, 2021), *Pantoea agglomerans* 33.1 (Quecine et al., 2012), and *Azospirillum brasilense* Ab-v5 (Hungria et al., 2010). Bmanagement constituted treatment with liquid Luria-Bertani medium only. Irrigation, weed control, and application of fertilizer (excluding nitrogen) were carried out according to crop requirements. The plants were evaluated when most had six expanded leaves, i.e., approximately 33 days after sowing. Manually measured phenotypes included plant height (PH), stalk diameter (SD), and shoot dry mass (SDM). Detailed information on the experimental design can be found in Yassue et al. (2021, 2022b).

### Genomic data

The 360 inbred lines were genotyped using the genotyping-by-sequencing method, followed by a two-enzyme (PstI and MseI) protocol (Sim et al., 2012; Poland et al., 2012). Deoxyribonucleic acid was extracted from leaves using cetyltrimethylammonium bromide (Doyle and Doyle, 1987). Single nucleotide polymorphism (SNP) calling was performed using TASSEL 5.0 (Bradbury et al., 2007) with B73 (B73-RefGen v4) as a reference genome. Single nucleotide polymorphism markers were removed if the call rate was *<* 90%, non-biallelic, or if the minor allele frequency was *<* 5%. Missing marker codes were imputed using Beagle 5.0 software (Browning et al., 2018). Markers with pairwise linkage disequilibrium *>* 0.99 were removed using the SNPRelate R package (Zheng et al., 2012). In total, 13,826 SNPs were retained after quality control. Detailed information on population genomics is available in Yassue et al. (2021).

### Hyperspectral imaging and processing

Leaves of B+ and B-plants were collected, and hyperspectral images were recorded using a benchtop Pika L. camera system (Resonon, Bozeman, MT, USA). The middle portion of the last completely expanded leaf was used as the region of interest for hyperspectral imaging. A dark room with an additional light supply was used to minimize light variation. Radiometric calibration was performed according to the manufacturer’s instructions. For each plant, a hyperspectral cube image was produced, which contained 150 bands with wavelengths in the 386–1,021 nm range. Image processing through the Spectral Python (Boggs, 2014) module was performed by applying a mask to remove the background from the image, and the mean reflectance of each pixel was used for further analysis. Hyperspectral imaging and processing details are described in Yassue et al. (2022a). In addition, 131 hyperspectral indices (mathematical band combinations) were calculated based on the mean reflectance value for each wavelength using the R package hsdar (Lehnert et al., 2019). These hyperspectral indices have been reported to be associated with a variety of phenotypes, such as nutrient and chlorophyll content, pigments, photosynthesis, and water content (Zarco-Tejada et al., 2004, 2005; Ranjan et al., 2012; Gitelson et al., 2014). A summary of the hyperspectral indices and their correlations are presented in Supplementary Tables S1–S5 and Supplementary Figure S1, respectively.

### Univariate BayesC

BayesC (Kizilkaya et al., 2010) was applied to estimate the markers effect and variance components using the following model.

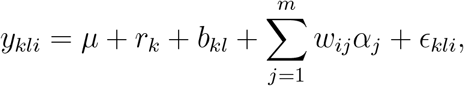

where *y*_*kli*_ is the vector of phenotypes (manually measured or hyperspectral phenotypes) for the *k*th repetition, *l*th block within repetition, and *i*th genotype; *µ* is the overall mean; *r* and *b* are the fixed effects for replication and block within replication, respectively; *w*_*ij*_ is the incidence matrices of marker covariates for each SNP coded as 0, 1, or 2; and *α*_*j*_ is the *j*th marker effect. The prior of *α*_*j*_ was:

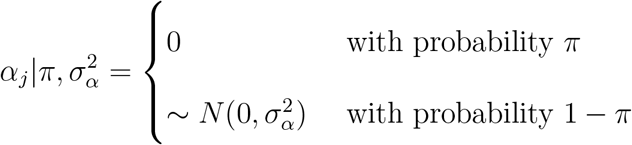

where 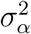 is the common marker genetic variance, *π* is a mixture proportion set to 0.99, and *ϵ* is the vector of residuals. A Gaussian prior 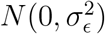 was assigned to the vector of residuals and a flat prior was assigned to *µ, r*, and *b*. The scaled inverse *χ*^2^ distribution was assigned to 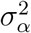 and 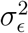 with the degrees of freedom equal to 4 and choosing the scale parameter such that the prior mean of the variance equals half of the phenotypic variance. The variance components obtained from univariate BayesC were used to estimate genomic heritability 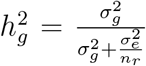. where 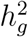 is the genomic heritability, 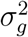 and 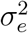 are the additive genomic and residual variances, respectively, and *n*_*r*_ is the number of replication (2).

### Bivariate BayesC

Bivariate BayesC was used to estimate the genetic correlation between manually measured and hyperspectral phenotypes. The model description follows that of univariate BayesC with some modification. Here, **y** is the vector of manually measured and hyperspectral phenotypes, and the marker effect of trait *t* for locus *j* followed

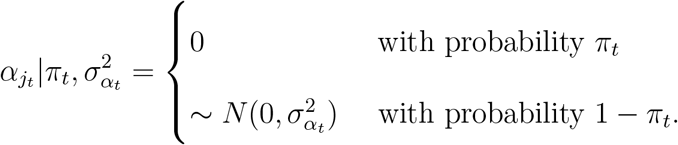

The *j*th marker effect can be reparameterized as ***α***_*j*_ = ***D***_*j*_***β***_*j*_, where ***D***_***j***_ is a diagonal matrix with elements *diag*(**D**_*j*_) = ***δ***_*j*_ = (*δ*_*j*1_, *δ*_*j*2_) indicating whether the *j*th marker effect for trait *t* is zero or non-zero, and ***β***_*j*_ follows a multivariate normal distribution with null mean and covariance matrix 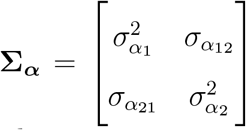, where *α, α*, and *α* (*α*) are marker genetic variance for trait 1, marker genetic variance for trait 2, and marker genetic covariance between traits 1 and 2, respectively, and the residuals were assumed independently and identically distributed multivariate normal vectors with null mean and covariance matrix **Σ**_***ϵ***_ (Cheng et al., ***2018b). The covariance matrices, Σ***_***α***_ and **Σ**_***ϵ***_, were assigned an inverse Wishart prior distribution with **W**^*−*1^(*S*_*α*_, *ν*_*α*_) and **W**^*−*1^(*S*_*ϵ*_, *ν*_*ϵ*_), respectively. We assumed all possible combinations for ***δ***_*j*_, namely, (0,0), (0,1), (1,0), and (1,1) having nonzero probability.

### Bayesian GWA analysis

The aforementioned BayesC was used to perform GWA analyses of hyperspectral reflectance values and manually measured phenotypes. Candidate markers were selected based on their posterior inclusion probabilities. The posterior inclusion probability indicates the probability of inclusion of a given marker in the model (Fernando and Garrick, 2013). According to a previous study (Fan et al., 2011), a posterior inclusion probability threshold of 0.10 was used for manually measured phenotypes, and 0.50 was used as a more conservative threshold for hyperspectral GWA. All Bayesian analyses were fit using 60,000 Markov chain Monte Carlo samples, 6,000 burn-ins, and a thinning rate of 60 implemented in JWAS (Cheng et al., 2018a). Model convergence was assessed using trace plots of the posterior means of the parameters. For each selected SNP associated with hyperspectral patterns, genes within an interval of 50 kilobase pair (kbp) upstream and downstream of the SNP were explored using the MaizeMine V1.3 server (Shamimuzzaman et al., 2020). One challenge was the many Manhattan plots that potentially needed to be generated owing to the three manually measured phenotypes, 150 hyperspectral wavelengths, and 131 hyperspectral indices. Instead of including all Manhattan plots as supplementary files, we developed a Shiny web application using the R package shiny (Chang et al., 2021), which provides functions for constructing interactive web applications. This application allows users to interactively explore all possible genome-to-phenome association combinations that are not elaborated on here.

### Results

### Estimates of genomic heritability and correlation

The genomic heritability estimates of manually measured PH, SD, and SDM were 0.61, 0.60, 0.30 in B+ plants and 0.57, 0.39, and 0.28 in B-plants, respectively. Plant height had the highest genomic heritability estimates, whereas SDM had the lowest. The means (and standard deviations) of genomic heritability estimates for hyperspectral wavelengths and indices were 0.45 (0.053) and 0.41 (0.081), respectively (Figure 1). The individual hyperspectral wavelengths that explained the largest genomic heritability estimates were 645 and 649 nm (B+), and 512 and 507 nm (B-), respectively. The estimates of genomic heritability showed a decreasing tendency as wavelength increased. Among the hyperspectral indices, NDNI and NPQI for B-and EVI for B+ showed the highest genomic heritability estimates. Although the mean of genomic heritability estimates of hyperspectral indices was lower than that of hyperspectral wavelengths, some indices explained more of the genetic variance than individual wavelengths. Overall, similar genomic heritability estimates were observed in B+ and B-plants for most wavelengths or indices.

**Figure 1:**
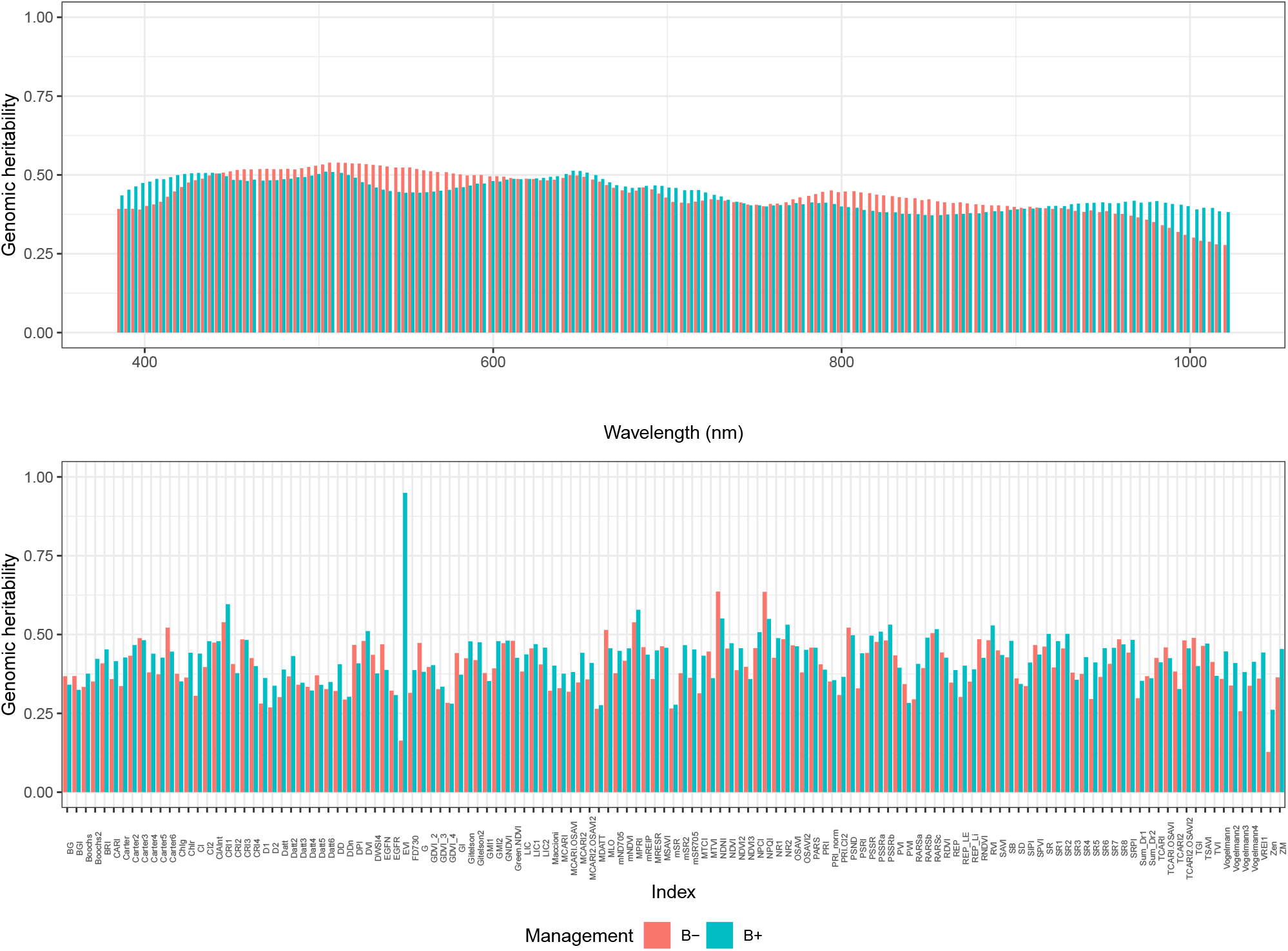
Genomic heritability for 150 hyperspectral reflectance values and 131 hyperspectral indices without (B-) and with (B+) plant growth-promoting bacteria inoculation.

The genomic correlation estimates between hyperspectral wavelengths and manually measured traits were largely positive (Figure 2). Genomic correlation estimates of PH ranged from -0.34 to 0.34. Positive correlations were observed for both B-and B+ at wavelengths from 400–700 nm, whereas negative correlations were observed only for B+ *>* 700 nm. In absolute terms, the hyperspectral wavelengths that provided the largest genomic correlation estimates were 578 nm (0.286) and 398 nm (0.193) for B-and B+, respectively. The extent of genomic correlation estimates was low in SD. The estimates were mostly positive, except for the beginning and end of wavelengths. The hyperspectral wavelengths showed positive correlations throughout the entire spectral range, except at the start of the SDM wavelength. In particular, higher correlations were observed at 700–1000 nm for B-. The hyperspectral wavelengths that provided the largest genomic correlation estimates with SDM were 817 nm (0.392) and 734 nm (0.235) for B-and B+, respectively. In contrast, genomic correlation estimates varied markedly across hyperspectral indices. The extent of the correlation estimates was lower regarding SD, compared to those of PH and SDM. The hyperspectral indices that provided the largest genomic correlation estimates with PH were D2 (0.339) and RDVI (−0.292) for B-and B+, respectively, whereas those with SDM were RARSb (−0.357) and NPQI (−0.340) for B-and B+, respectively.

**Figure 2:**
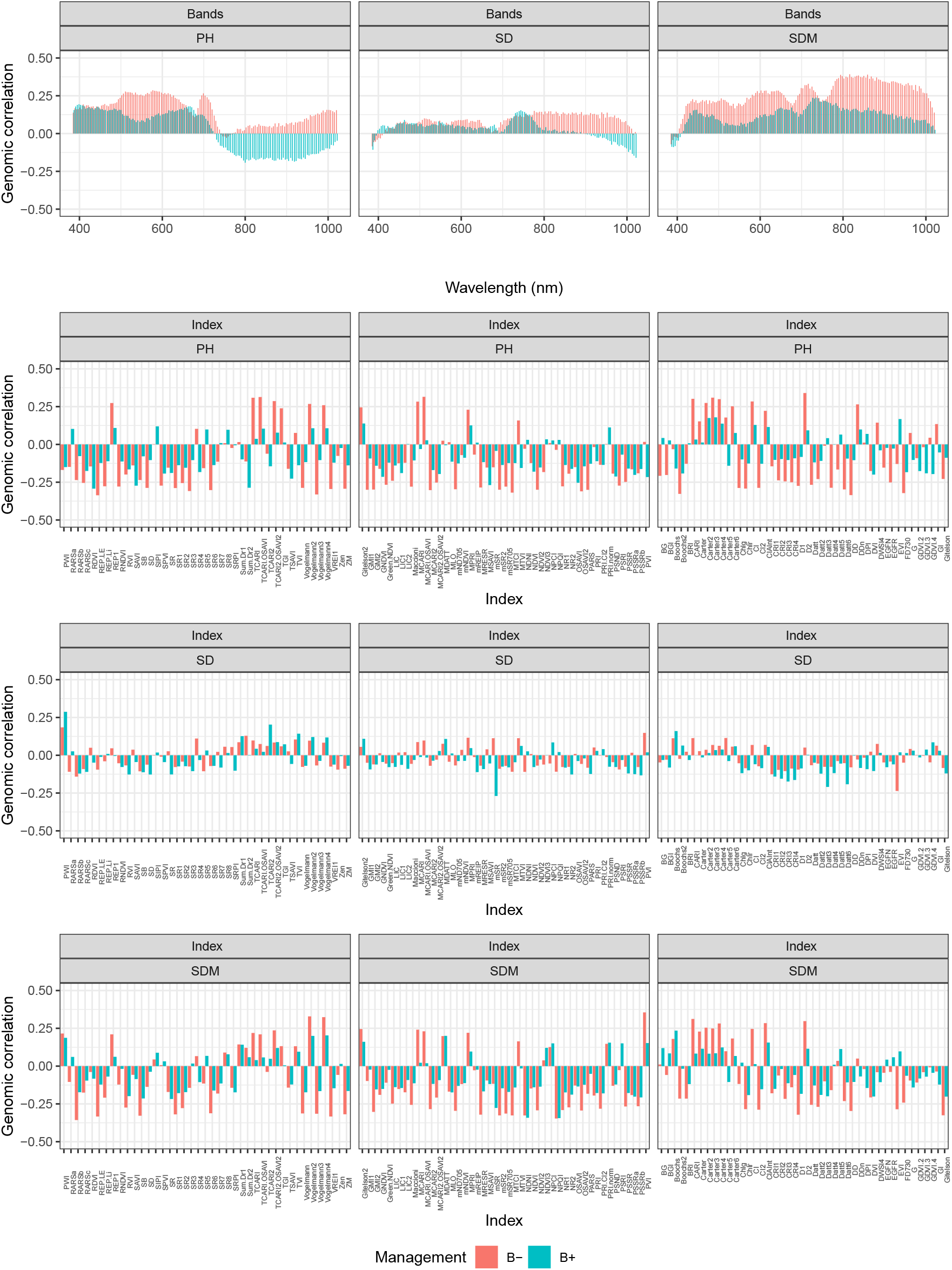
Genomic correlations between manually measured phenotypes and hyperspectral reflectance values and hyperspectral indices under without (B-) and with (B+) plant growthpromoting bacteria inoculation.

### GWA analyses for growth-related and hyperspectral traits

A total of 86 SNPs were selected from the BayesC analysis using the posterior inclusion probability threshold of 0.10 for PH, SD, and SDM (Figure 3 and Supplementary Tables S6–S10). Plant height showed the highest number of selected markers (21 and 24 for B+ and B-, respectively), whereas SDM had the lowest (5 for B+); no SDM markers were detected in B-. No overlapping SNPs were identified across manually measured phenotypes, whereas only four SNPs were selected for both PGPB inoculation conditions (B-and B+), indicating that PGPB may alter plant growth patterns and the genomic regions controlling them. A conservative posterior inclusion probability threshold of 0.50 was used to find SNPs associated with the hyperspectral-derived phenotypes and candidate genes. Of the 25 detected SNPs, five SNPs were associated with at least five different hyperspectral phenotypes. Gene annotation of each selected SNP within an interval of 100 kbp showed the presence of genes that have been previously reported as related to growth-related phenotypes or responses to abiotic stressors (Table 1). The hyperspectral indices Chlg, CRI2, CRI3, CRI4, Datt6, GMI1, PARS, SD, and SR3 were associated with genes nrt2, nrt2.2, and Zm00001d054060 on chromosome 4. The index CRI1 was associated with gene Zm00001d012924 on chromosome 5, and the EVI index was associated with genes Zm00001d029820 and Zm00001d007843 on chromosomes 1 and 2, respectively, and 100275163 on chromosome 6. In addition, hyperspectral wavelengths from 398 to 434 nm were associated with gene Zm00001d012719 on chromosome 8.

**Figure 3:**
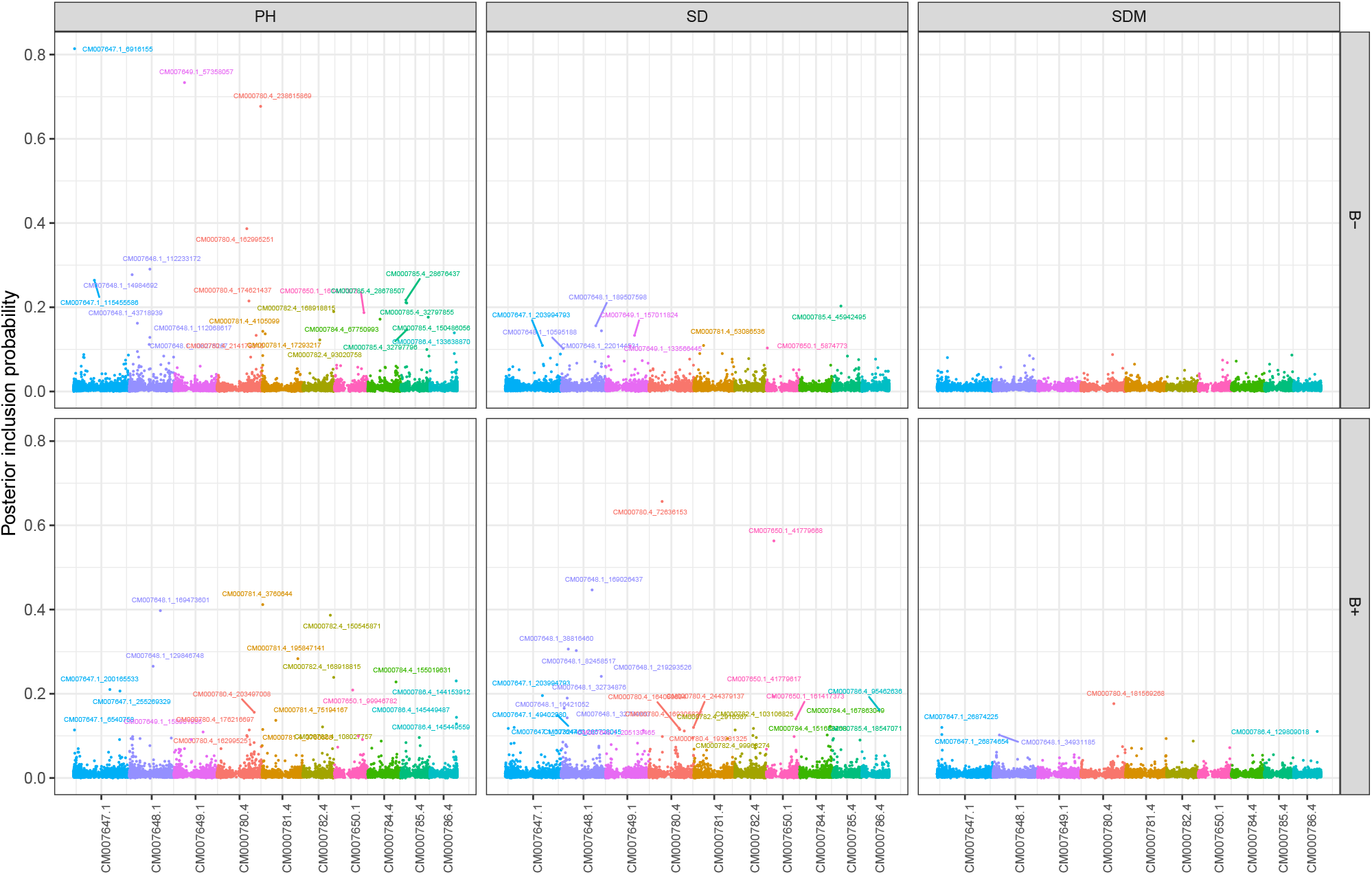
Genome-wide association analysis of manually measured phenotypes without (B-) and with (B+) plant growth-promoting bacteria inoculation. Plant height (PH); stalk diameter (SD) and shoot dry mass (SDM).

**Table 1:** Selected single nucleotide polymorphisms markers based on BayesC using the posterior inclusion probability threshold of 0.50 for 281 hyperspectral phenotypes under with (B+) or without (B-) plant growth-promoting bacteria inoculation.

| Management | Chr<sup>1</sup> | Marker ID<sup>2</sup> | Pheno<sup>3</sup> | MAF<sup>4</sup> | PIP<sup>5</sup> | NG<sup>6</sup> | Candidate gene |
| --- | --- | --- | --- | --- | --- | --- | --- |
| B- | 4 | CM000780.4_164335549 | 1 | 0.27 | 0.508 | 2 |  |
| B- | 8 | CM000784.4_174418506 | 5 | 0.17 | 0.551 | 9 |  |
| B+ | 1 | CM007647.1_69298773 | 1 | 0.06 | 0.981 | 6 |  |
| B+ | 1 | CM007647.1_80258452 | 1 | 0.36 | 0.820 | 3 |  |
| B+ | 1 | CM007647.1_88464412 | 1 | 0.18 | 0.661 | 7 | Zm00001d029820 |
| B+ | 1 | CM007647.1_113763491 | 1 | 0.13 | 0.894 | 5 |  |
| B+ | 1 | CM007647.1_173938755 | 1 | 0.06 | 0.966 | 5 |  |
| B+ | 2 | CM007648.1_181732219 | 1 | 0.27 | 0.549 | 8 |  |
| B+ | 2 | CM007648.1_240687544 | 1 | 0.08 | 0.851 | 4 | Zm00001d007843 |
| B+ | 2 | CM007648.1_214786010 | 14 | 0.46 | 0.602 | 9 |  |
| B+ | 4 | CM000780.4_245633076 | 9 | 0.38 | 0.585 | 22 | nrt2, nrt2.2<br>Zm00001d054060 |
| B+ | 4 | CM000780.4_206259028 | 1 | 0.06 | 0.559 | 8 |  |
| B+ | 5 | CM000781.4_2005858 | 1 | 0.32 | 0.581 | 15 | Zm00001d012924 |
| B+ | 5 | CM000781.4_213988333 | 1 | 0.33 | 0.604 | 7 |  |
| B+ | 6 | CM000782.4_108027757 | 5 | 0.48 | 0.519 | 10 |  |
| B+ | 6 | CM000782.4_9906627 | 1 | 0.07 | 0.633 | 12 | 100275163, 100192849 |
| B+ | 7 | CM007650.1_22239405 | 1 | 0.15 | 0.570 | 2 |  |
| B+ | 8 | CM000784.4_177771765 | 1 | 0.05 | 0.826 | 13 |  |
| B+ | 8 | CM000784.4_179214567 | 10 | 0.48 | 0.593 | 11 | Zm00001d012719 |
| B+ | 9 | CM000785.4_127613348 | 1 | 0.32 | 0.510 | 3 |  |
| B+ | 10 | CM000786.4_88567864 | 1 | 0.10 | 0.954 | 6 |  |
| B+ | 10 | CM000786.4_118526736 | 1 | 0.27 | 0.569 | 3 |  |
| B+ | 10 | CM000786.4_122822696 | 1 | 0.13 | 0.969 | 12 |  |
| B+ | 10 | CM000786.4_137486546 | 1 | 0.08 | 0.996 | 9 |  |
| B+ | 10 | CM000786.4_145821750 | 1 | 0.16 | 0.544 | 6 |  |

<sup>1</sup> Chromosome number
<sup>2</sup> Each marker ID is comprised of chromosome ID and marker location that are separated by the underscore sign.
<sup>3</sup> The number of phenotypes influenced by the marker.
<sup>4</sup> Minor allele frequency.
<sup>5</sup> Average of posterior inclusion probability for the selected phenotypes
<sup>6</sup> Number of genes within the gene interval.

### Integration of GWA analyses

Eight SNPs influenced both the manually measured and hyperspectral phenotypes, which were visualized in a phenome-wide association plot (Figure 4). In general, most SNPs were identified in B+ plants. Two SNPs, CM000780.4 181569268 and CM000782.4 108027757, exhibited a strong association with a wide range of hyperspectral phenotypes, along with PH and SDM in B+ management. A list of candidate genes influencing both manually measured and hyperspectral phenotypes revealed that they may play an important role in nitrogen uptake and plant responses to biotic and abiotic stressors (Table 2). A Shiny web application was developed to explore the multi-phenotype genome-wide association results interactively (https://github.com/vt-ads/ShinyGWASPheWAS).

**Figure 4:**
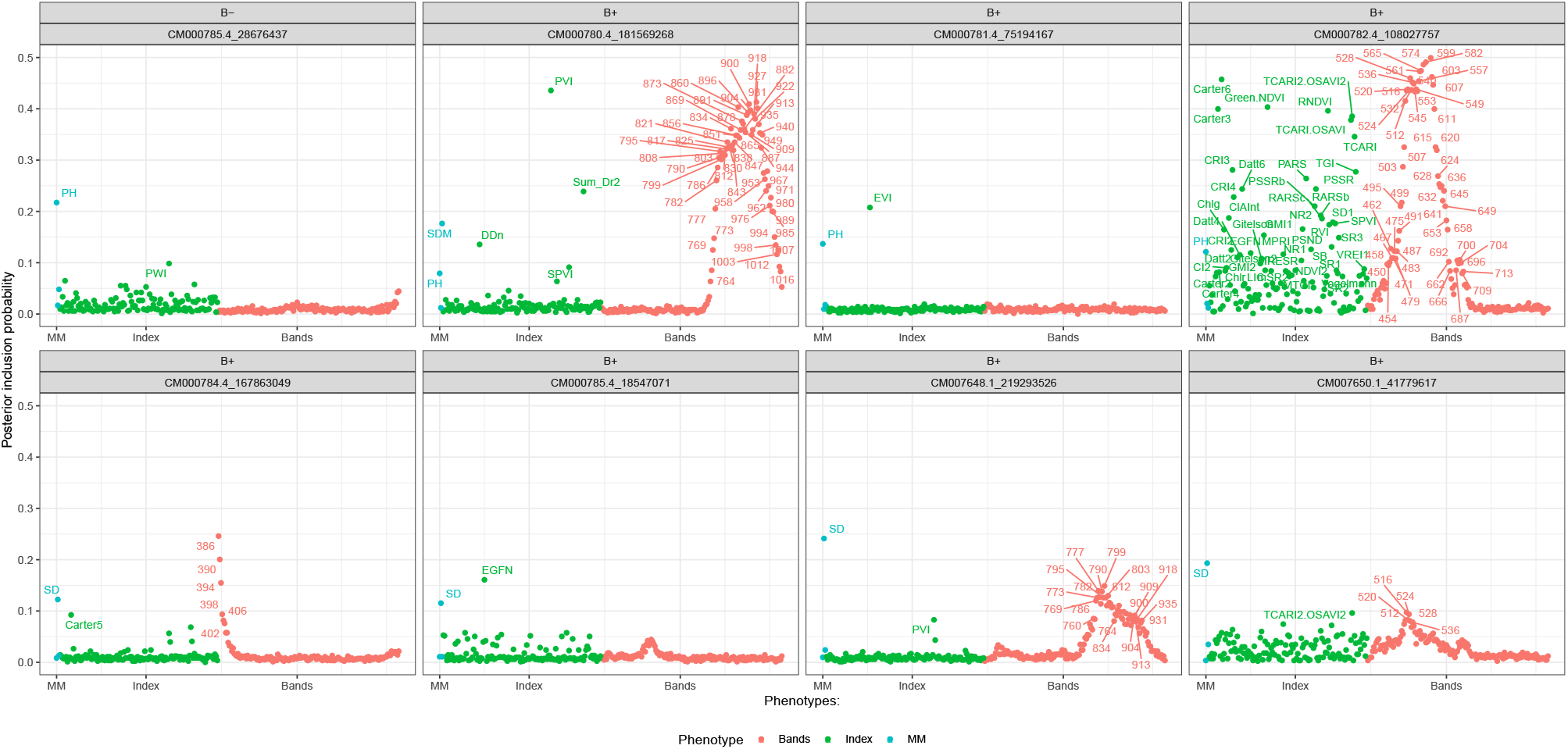
Phenome-wide association analysis plot of selected eight markers having influence on both manually measured and hyperspectral phenotypes without (B-) and with (B+) plant growth-promoting bacteria inoculation. Plant height (PH); stalk diameter (SD); shoot dry mass (SDM); hyperspectral bands (B-ands); hyperspectral index (Index); and manually measured (MM). The abbreviations of hyperspectral indices are defined in Tables S1-S4.

**Table 2:**
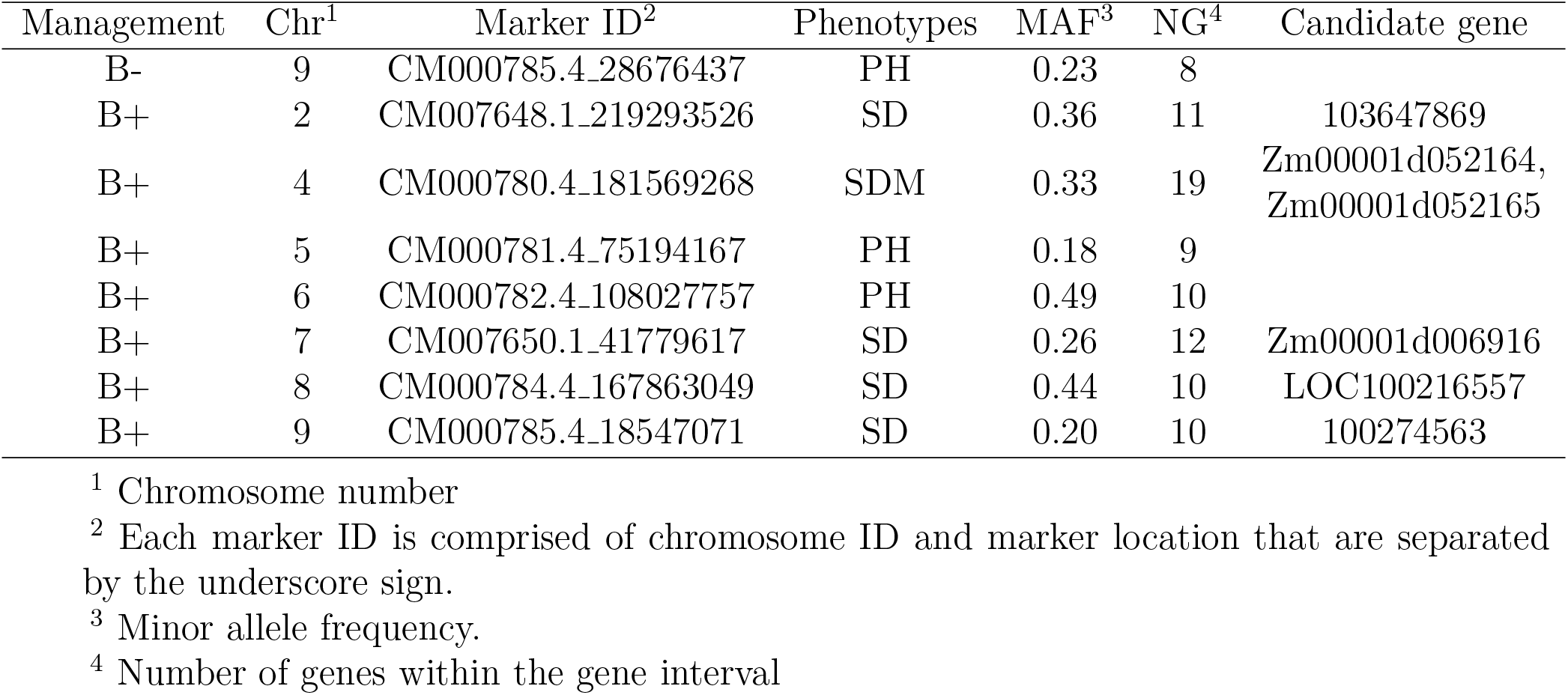
List of candidate genes influencing both manually measured phenotypes and hyperspectral phenotypes. Selected single nucleotide polymorphisms markers and their candidate genes influencing both manually measured phenotypes and hyperspectral phenotypes under with (B+) or without (B-) plant growth-promoting bacteria inoculation.

## Discussion

The utility of hyperspectral imaging technology for phenotyping has recently gained increasing attention because such data can capture the resonance of certain physicochemical compounds in plants. Growth-related traits, such as PH, SD, and SDM, are complex traits controlled by many genes with small individual effects. Therefore, we expected that hyperspectral imagery-based data would shed light on the underlying genetic factors and help assess the variability of maize to aid in identifying candidate genes. However, translating hyperspectral reflectance values or indices into a biological context, such as metabolic, morphological, or functional changes, can be difficult and time-consuming. This study used GWA analysis of manually measured phenotypes, single-band reflectance, and hyperspectral indices were used to investigate the genetic basis of responses to PGPB. BayesC, which performs variable selection, was applied for GWA analysis. The posterior inclusion probability of each marker was used to identify the relevant SNPs. The preference for using posterior inclusion probability instead of window posterior probability of association (Fernando et al., 2017) was due to the low marker density and unequal distribution of SNPs across the genome in the maize population.

### Estimates of genomic heritability and correlation

The hyperspectral phenotypes showed a similar range of genomic heritability estimates relative to that of manually measured phenotypes. For most hyperspectral-derived phenotypes, the heritability estimates varied from 0.30 to 0.50, indicating that hyperspectral data can capture genetic variation. In addition, the genetic correlation between the manually measured and hyperspectral phenotypes indicated that the same sets of genes probably influenced these responses.

The relatively higher genomic correlation estimates for PH in the spectral range of 400–700 nm (visible spectrum) of B-plants may indicate an association between plant height and leaf pigments, such as carotenoids, chlorophyll a and b, and nitrogen concentrations (Zhao et al., 2003; Ayala-Silva and Beyl, 2005). Similarly, higher genomic correlation estimates were observed for SDM at 700–1,000 nm (near-infrared). The association between near-infrared spectra and plant biomass in maize was reported in maize in a previous study (Ma et al., 2020). Wavelengths in this range have also been reported to indicate nitrogen content in rapeseed (Müller et al., 2008) and wheat (Hansen and Schjoerring, 2003).

### Genome-wide association analysis

Overall, more SNP associations were observed in B+ than in B-plants. However, the genomic heritability estimates were similar between the two managements. This suggests that the genetics underlying hyperspectral responses differ between the two managements. Genome-wide association analysis showed that SNP CM000780.4 245633076 was associated with nine hyperspectral phenotypes pointing to three candidate genes (nrt2, nrt2.2, and Zm00001d054060). These genes have been previously reported as part of 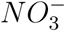 transporter gene families and as candidates for nitrate uptake along the primary maize root (Liu et al., 2009; Sorgonà et al., 2011; Wang et al., 2020). The genes Zm00001d029820 and Zm00001d012924 are involved in plant development and environmental stress conditions (Zhang et al., 2020; Zhu et al., 2021). The candidate gene Zm00001d007843 may affect kernel size (Zhou et al., 2021), and gene Zm00001d012719 is a candidate transcription factor mediating plant responses to abiotic stressors (Vendramin et al., 2020). We did not directly evaluate the phenotypes related to previously reported candidate genes. Nevertheless, hyperspectral signatures may be of use for indirectly assessing these phenotypes. Furthermore, eight SNPs were detected in both the manually measured and hyperspectral phenotypes. Genes Zm00001d052164 and Zm00001d052165 on chromosome 4 regulate nitrogen assimilation (Wang et al., 2020), and Zm00001d006916 is a candidate gene responsible for autophagy, which may play a role regarding responses to abiotic stressors (Tang and Bassham, 2018). The LOC100216557 gene is associated with resilience of maize to aphids and may be responsible for plant defense responses and stress tolerance (Srivastava et al., 2018), and 103647869 is a candidate gene for resistance to *Aspergillus flavus* infection or aflatoxin contamination (Liu et al., 2021). The gene 100274563 is associated with ear weight per plant (Zhou et al., 2020).

The Shiny web application with an interactive interface is a powerful tool for the visualization and interpretation of GWA analysis of multiple phenotypes. Two-way Manhattan plots can be used to investigate all associations across traits and managements, including non-significant results that are not elaborated on here. In addition, phenome-wide association plots can be used to identify and visualize markers with mutual influence across hundreds or thousands of phenotypes. The Shiny application can be easily extended to other high-throughput phenotyping data, such as longitudinal, fluorescence, and thermal data.

## Conclusions

The hyperspectral signatures captured the genetic variability in the maize diversity panel and were associated with growth-related traits under PGPB inoculation. Genome-wide association analysis of hyperspectral data identified genomic regions that influenced both manually measured phenotypes and hyperspectral bands. In addition, a Shiny web application for multiple-phenotype GWA was developed.

## Abbreviations

(GWA): genome-wide association
(PGPB): plant growth-promoting bacteria
(PH): plant height
(SDM): shoot dry mass
(SNPs): single-nucleotide polymorphisms
(SD): stalk diameter
(B+): with plant growth-promoting bacterial inoculation
(B-): and without plant growth-promoting bacterial inoculation

## Acknowledgments

The authors acknowledge Pedro Takao Yamamoto and Fernando Henrique Iost Filho for their support in collecting the hyperspectral images.

## Author contributions

Rafael Massahiro Yassue: Conceptualization; Data curation, Formal analysis, Investigation; Methodology; Visualization; Writing-original draft; Writing-review & editing. Giovanni Galli: Investigation; Methodology; Writing-review & editing. Chun-Peng James Chen: Methodology; Writing-review & editing. Roberto Fritsche-Neto: Conceptualization; Funding acquisition; Supervision; Writing-review & editing. Gota Morota: Conceptualization; Methodology; Funding acquisition; Supervision; Writing-original draft; Writing-review & editing.

