## Supplemental material for "Genome-wide association analysis of hyperspectral reflectance data to dissect growth-related traits genetic architecture in maize under inoculation with plant growth-promoting bacteria"

### Supplementary Tables

Table S1: List of hyperspectral indices used in the study.

| Index | Abbreviation | Formula | Reference |
|---|---|---|---|
| Boochs index | Boochs | $D_{703}$ | (Boochs et al., 1990) |
| Boochs index 2 | Boochs2 | $D_{720}$ | (Boochs et al., 1990) |
| Carter index | Carter | $R_{695}/R_{420}$ | (Carter, 1994) |
| Carter index 2 | Carter2 | $R_{695}/R_{760}$ | (Carter, 1994) |
| Carter index 3 | Carter3 | $R_{605}/R_{760}$ | (Carter, 1994) |
| Carter index 4 | Carter4 | $R_{710}/R_{760}$ | (Carter, 1994) |
| Carter index 5 | Carter5 | $R_{695}/R_{670}$ | (Carter, 1994) |
| Carter index 5 | Carter6 | $R_{550}$ | (Carter, 1994) |
| Chlorophyll index | CI | $R_{675} * R_{690}/R_{683}^2$ | (Zarco-Tejada et al., 2013) |
| Chlorophyll index 2 | CI2 | $R_{760}/R_{700} - 1$ | (Gitelson et al., 2003) |
| Chlorophyll absorption integral | ClAInt | $\int_{60n\ \text{nm}}^{735\ \text{nm}} \text{R}$ | (Oppelt and Mauser, 2004) |
| Carotenoid reflectance index 1 | CRI1 | $1/R_{515} - 1/R_{550}$ | (Gitelson et al., 2003) |
| Carotenoid reflectance Iindex 2 | CRI2 | $1/R_{515} - 1/R_{770}$ | (Gitelson et al., 2003) |
| Carotenoid reflectance index 3 | CRI3 | $1/R_{515} - 1/R_{550} * R_{770}$ | (Gitelson et al., 2003) |
| Carotenoid reflectance index 4 | CRI4 | $1/R_{515} - 1/R_{700} * R_{770}$ | (Gitelson et al., 2003) |
| Simple Ratio 730/706 | D1 | $D_{730}/D_{706}$ | (Zarco-Tejada et al., 2013) |
| Simple Ratio 705/722 | D2 | $D_{705}/D_{722}$ | (Zarco-Tejada et al., 2013) |
| Datt index | Datt | $(R_{850} - R_{710})/(R_{850} - R_{680})$ | (Datt, 1999) |
| Datt index2 | Datt2 | $R_{850}/R_{710}$ | (Datt, 1999) |
| Datt index 3 | Datt3 | $D_{754}/D_{704}$ | (Datt, 1999) |
| Datt index 4 | Datt4 | $R_{672}/(R_{550} * R_{708})$ | (Datt, 1999) |
| Datt index 5 | Datt5 | $R_{672}/R_{550}$ | (Datt, 1999) |
| Datt index 6 | Datt6 | $(R_{860})/(R_{550} * R_{708})$ | (Datt, 1999) |
| Double difference index | DD | $(R_{749} - R_{720}) - (R_{701} - R_{672})$ | (le Maire et al., 2004) |
| New double difference index | DDn | $2 * (R_{710} - R_{660} - R_{760})$ | (le Maire et al., 2008) |
| Double peak index | DPI | $(D_{688} * D_{710})/D_{697}^2$ | (Zarco-Tejada et al., 2013) |
| Disease water stress index | DWSI4 | $R_{550}/R_{680}$ | (Apan et al., 2004) |
| Normalized ratio between the maxima of the first derivatives of reflectances at the red edge and green regions | EGFN | $(\max(D_{650:750}) - \max(D_{500:550}))/(\max(D_{650:750}) + \max(D_{500:550}))$ | (Peñuelas et al., 1994) |
| Ratio between dRE and dG | EGFR | $\max(D_{650:750})/\max(D_{500:550})$ | (Peñuelas et al., 1994) |
| Enhanced vegetation index | EVI | $25 * ((R_{800} - R_{670})/(R_{800} - (6 * R_{670}) - (75 * R_{475}) + 1))$ | (Huete et al., 1997) |
| Generalized difference vegetation index | GDVI_2 | $(R_{800}^n - R_{680}^n)/(R_{800}^n + R_{680}^n)$ | (Wu, 2014) |
| Generalized difference vegetation index | GDVI_3 | | (Wu, 2014) |
| Generalized difference vegetation index | GDVI_4 | | (Wu, 2014) |
| Greenness index | GI | $R_{554}/R_{677}$ | (Smith et al., 1995) |
| Gitelson index | Gitelson | $1/R_{700}$ | (Gitelson et al., 1999) |
| Gitelson index 2 | Gitelson2 | $(R_{750} - R_{800}/R_{695} - R_{740}) - 1$ | (Gitelson et al., 2003) |

Rxxx: Reflectance at wavelength 'xxx'.
Dxxx: First derivation of reflectance values at wavelength 'xxx'.
maxxxx: maximum value at wavelength 'xxx'.

Table S2: List of hyperspectral indices used in the study.

| Index | Abbreviation | Formula | Reference |
|---|---|---|---|
| Maccioni index | Maccioni | $(R_{780} - R_{710})/(R_{780} - R_{680})$ | (Maccioni et al., 2001) |
| Modified chlorophyll absorption in reflectance index | MCARI | $((R_{700} - R_{670}) - 0.2 * (R_{700} - R_{550})) * (R_{700}/R_{670})$ | (Daughtry et al., 2000) |
| | MCARI/OSAVI | | (Daughtry et al., 2000) |
| Modified chlorophyll absorption in reflectance index 2 | MCARI2 | $((R_{750} - R_{705}) - 0.2 * (R_{750} - R_{550})) * (R_{750}/R_{705})$ | (Wu et al., 2008) |
| | MCARI2/OSAVI2 | | (Wu et al., 2008) |
| Modified normalized difference at 705 nm wavelength | mND705 | $(R_{750} - R_{705})/(R_{750} + R_{705} - 2 * R_{445})$ | (Sims and Gamon, 2002) |
| Modified normalized difference vegetation index | mNDVI | $(R_{800} - R_{680})/(R_{800} + R_{680} - 2 * R_{445})$ | (Sims and Gamon, 2002) |
| Normalized difference physiological reflectance index | MPRI | $(R_{515} - R_{530})/(R_{515} + R_{530})$ | (Hernández-Clemente et al., 2011) |
| Modified red-edge inflection point | mREIP | red-edge inflection point using Gaussain fit | (MILLER et al., 1990) |
| Modified soil-adjusted vegetation index | MSAVI | $05 * (2 * R_{800} + 1 - ((2 * R_{800} + 1)^2 - 8 * (R_{800} - R_{670}))^{05})$ | (Qi et al., 1994) |
| Modified simple ratio of reflectance | mSR | $(R_{800} - R_{445})/(R_{680} - R_{445})$ | (Sims and Gamon, 2002) |
| Modified simple ratio of reflectance 2 | mSR2 | $(R_{750}/R_{705}) - 1/(R_{750}/R_{705} + 1)^{05}$ | (Chen, 1996) |
| Modified simple ratio of reflectance 705 | mSR705 | $(R_{750} - R_{445})/(R_{705} - R_{445})$ | (Sims and Gamon, 2002) |
| MERIS terrestrial chlorophyll index | MTCI | $(R_{754} - R_{709})/(R_{709} - R_{681})$ | (Dash and Curran, 2004) |
| Modified triangular vegetation index | MTVI | $1.2 * (1.2 * (R_{800} - R_{550}) - 25 * (R_{670} - R_{550}))$ | (Haboudane et al., 2002) |
| Chlorophyll absorption ratio index | Cari | | (Kim et al., 1994) |
| Normalized difference vegetation index | NDVI | $(R_{800} - R_{680})/(R_{800} + R_{680})$ | (Tucker, 1979) |
| Normalized difference vegetation index 2 | NDVI2 | $(R_{750} - R_{705})/(R_{750} + R_{705})$ | (Gitelson and Merzlyak, 1994) |
| Normalized difference vegetation index 3 | NDVI3 | $(R_{682} - R_{553})/(R_{682} + R_{553})$ | (Gandia et al., 2004) |
| Normalized pigment chlorophyll index | NPCI | $(R_{680} - R_{430})/(R_{680} + R_{430})$ | (Peñuelas et al., 1994) |
| Optimized soil-adjusted vegetation index | OSAVI | $(1+0.16)*(R_{800}-R_{670})/(R_{800}+R_{670}+0.16)$ | (Rondeaux et al., 1996) |
| Optimized soil-adjusted vegetation index 2 | OSAVI2 | $(1+0.16)*(R_{750}-R_{705})/(R_{750}+R_{705}+0.16)$ | (Wu et al., 2008) |
| Ratio analysis of reflectance spectra | PARS | $R_{746}/R_{513}$ | (Chappelle et al., 1992) |
| Photochemical reflectance index | PRI | $(R_{531} - R_{570})/(R_{531} + R_{570})$ | (Gamon et al., 1992) |
| Photochemical reflectance index normalized | PRI_norm | $PRI*(-1)/(RDVI*R_{700}/R_{670})$ | (Zarco-Tejada et al., 2013) |
| | PRI * CI2 | $PRI * CI2$ | (Garrity et al., 2011) |
| Plant senescence reflectance index | PSRI | $(R_{678} - R_{500}/R_{750}$ | (Merzlyak et al., 1999) |

Rxxx: Reflectance at wavelength 'xxx'.
Dxxx: First derivation of reflectance values at wavelength 'xxx'.
maxxxx: maximum value at wavelength 'xxx'.

Table S3: List of hyperspectral indices used in the study.

| Index | Abbreviation | Formula | Reference |
|---|---|---|---|
| Red-edge position linear interpolation | REP_LE | Red-edge position through linear extrapolation | (Cho and Skidmore, 2006) |
| Red-edge position linear interpolation | REP_Li | $R_{re} = (R_{670} + R_{780})/2$ | (Guyot and Baret, 1988) |
| Soil adjusted vegetation index | SAVI | $(1 + L) * (R_{800} - R_{670})/(R_{800} + R_{670} + L)$ | (Huete, 1988) |
| Structure insensitive pigment index | SIPI | $(R_{800} - R_{445})/(R_{800} - R_{680})$ | (Peñuelas et al., 1995; PEN¯UELAS et al., 1995) |
| spectral polygon vegetation index | SPVI | $0.4 * 3.7 * (R_{800} - R_{670}) - 1.2 * ((R_{530} - R_{670})^2)^{0.5}$ | (Vincini et al., 2006) |
| Simple ratio | SR | $R_{800}/R_{680}$ | (Jordan, 1969) |
| Simple ratio 1 | SR1 | $R_{750}/R_{700}$ | (Gitelson and Merzlyak, 1997) |
| Simple ratio 2 | SR2 | $R_{752}/R_{690}$ | (Gitelson and Merzlyak, 1997) |
| Simple ratio 3 | SR3 | $R_{750}/R_{550}$ | (Gitelson and Merzlyak, 1997) |
| Simple ratio 4 | SR4 | $R_{700}/R_{670}$ | (McMurtrey et al., 1994) |
| Simple ratio 5 | SR5 | $R_{675}/R_{700}$ | (Chappelle et al., 1992) |
| Simple ratio 6 | SR6 | $R_{750}/R_{710}$ | (Zarco-Tejada and Miller, 1999) |
| Simple ratio 7 | SR7 | $R_{440}/R_{690}$ | (Lichtenthaler et al., 1996) |
| Simple ratio 8 | SR8 | $R_{515}/R_{550}$ | (Hernández-Clemente et al., 2012) |
| Simple ratio pigment index | SRPI | $R_{430}/R_{680}$ | (PEN¯UELAS et al., 1995) |
| Sum of first derivative reflectance 1 | Sum_Dr1 | $\Sigma_{i=626}^{795} D1_i$ | (Elvidge and Chen, 1995) |
| Sum of first derivative reflectance 2 | Sum_Dr2 | $\Sigma_{i=680}^{780} D1_i$ | (Filella and Peñuelas, 1994) |
| Transformed chlorophyll absorption reflectance index | TCARI | $3 * ((R_{700} - R_{670}) - 02 * R_{700} - R_{550}) * (R_{700}/R_{670}))$ | (Haboudane et al., 2002) |
| | TCARI/OSAVI | $TCARI/OSAVI$ | (Haboudane et al., 2002) |
| Transformed chlorophyll absorption reflectance index 2 | TCARI2 | $3 * ((R_{750} - R_{705}) - 02 * (R_{750} - R_{550}) * (R_{750}/R_{705}))$ | (Wu et al., 2008) |
| Triangular greenness index | TGI | $-0.5(190(R_{670} - R_{550}) - 1.20(R_{670} - R_{480}))$ | (Hunt et al., 2013) |
| Triangular vegetation index | TVI | $0.5 * (120 * (R_{750} - R_{550}) - 200 * (R_{670} - R_{550}))$ | (Broge and Leblanc, 2001) |
| Vogelmann index | Vogelmann | $R_{740}/R_{720}$ | Vogelmann et al. (1993) |
| Vogelmann index 2 | Vogelmann2 | $(R_{734} - R_{747})/(R_{715} + R_{726})$ | Vogelmann et al. (1993) |
| Vogelmann index 3 | Vogelmann3 | $D_{715}/D_{705}$ | Vogelmann et al. (1993) |
| Vogelmann index 4 | Vogelmann4 | $(R_{734} - R_{747})/(R_{715} + R_{720})$ | Vogelmann et al. (1993) |
| Ratio vegetation index | RVI | $R790/R650$ | (Pearson and Miller, 1972) |

Rxxx: Reflectance at wavelength 'xxx'.
Dxxx: First derivation of reflectance values at wavelength 'xxx'.
maxxxx: maximum value at wavelength 'xxx'.

Table S4: List of hyperspectral indices used in the study.

| Index | Abbreviation | Formula | Reference |
|---|---|---|---|
| Red-edge position linear interpolation | REP | $R_{700} + 40 * ((R_{670} + R_{780})/2 - R_{700})/(R_{740} - R_{700}))$ | (Ali and Imran, 2020) |
| First derivative | FD730 | $D_{730}$ | (Peng et al., 2018) |
| Modified Datt index | MDATT | $(R_{719} - R_{726})/(R_{719} - R_{743})$ | (Velichkova and Krezhova, 2019) |
| Modified red-edge simple ratio | MRESR | $(R_{750} - R_{450})/(R_{705} + R_{450})$ | (Velichkova and Krezhova, 2019) |
| Near infrared/Red 1 | NR1 | $R_{760}/R_{695}$ | (Velichkova and Krezhova, 2019) |
| Near infrared/Red 2 | NR2 | $R_{800}/R_{650}$ | (Velichkova and Krezhova, 2019) |
| Zarco - Miller Index | ZM | $R_{750}/R_{710}$ | (Velichkova and Krezhova, 2019) |
| Greenness | G | $R_{554}/R_{667}$ | (Velichkova and Krezhova, 2019) |
| Blue/Green | BG | $R_{677}/R_{554}$ | (Velichkova and Krezhova, 2019) |
| Vogelmann red edge | VREI1 | $R_{740}/R_{720}$ | (Velichkova and Krezhova, 2019) |
| Single band index | SB | $1/R_{700}$ | (Velichkova and Krezhova, 2019) |
| Single-difference index | SD | $(1/R_{550}) - (1/R_{750})$ | (Velichkova and Krezhova, 2019) |
| Green Chl index | Chlg | $R_{760}/R_{550} - 1$ | (Velichkova and Krezhova, 2019) |
| Red edge Chl index | Chlr | $R_{760}/R_{714} - 1$ | (Velichkova and Krezhova, 2019) |
| Simple ratio pigment specific simple ratio (Cholophyll a) | PSSRa | $R_{800}/R_{675}$ | (Blackburn, 1998a) |
| Simple ratio pigment specific simple ratio B1 | PSSRb | $R_{800}/R_{650}$ | (Blackburn, 1998a) |
| Normalized pheophytinization index | NPQI | $(R_{415} - R_{435})/(R_{415} + R_{435})$ | (PEN¯UELAS et al., 1995) |
| Normalized difference nitrogenindex | NDNI | $(R_{415} - R_{435})/(R_{415} + R_{435})$ | (Wang and Wei, 2016) |
| | MLO | $R_{531}/R_{645}$ | (Meiforth et al., 2020) |
| Lichtenthaler index | LIC | $(R_{850} - R_{710})/(R_{850} + R_{680})$ | (Lichtenthaler et al., 1996) |
| Lichtenthaler index 1 | LIC1 | $(R_{800} - R_{680})/(R_{800} + R_{680})$ | (Lichtenthaler et al., 1996) |
| Lichtenthaler index 2 | LIC2 | $R_{440}/R_{690}$ | (Lichtenthaler et al., 1996) |
| Blue/Green pigment index | BGI | $R_{450}/R_{550}$ | (ZARCOTEJADA et al., 2005) |
| Blue/Red pigment index | BRI | $(R_{450}/R_{690})$ | (ZARCOTEJADA et al., 2005) |
| Ratio analysis of reflectance spectra chlorophyll a | RARSa | $(R_{675}/R_{700})$ | (Kycko et al., 2019) |
| Ratio analysis of reflectance spectra chlorophyll b | RARSb | $(R_{675}/(R_{650} * R_{700}))$ | (Montesinos-López et al., 2017) |

Rxxx: Reflectance at wavelength 'xxx'.
Dxxx: First derivation of reflectance values at wavelength 'xxx'.
maxxxx: maximum value at wavelength 'xxx'.

Table S5: List of hyperspectral indices used in the study.

| Index | Abbreviation | Formula | Reference |
| --- | --- | --- | --- |
| Gitelson and Merzylak index 1 | GMI1 | $R_{750}/R_{550}$ | (Gitelson et al., 2003) |
| Gitelson and Merzylak index2 | GMI2 | $R_{750}/R_{700}$ | (Gitelson et al., 2003) |
| Green normalized difference vegetation indexx | GreenNDVI | $(R_{800} - R_{550})/(R_{800} + R_{550})$ | (Gitelson et al., 1996) |
| Simple ratio 800/635 pigment specific simple ratio (Cholophyll b) | PSSR | $R_{800}/R_{635}$ | (Blackburn, 1998b) |
| Pigment specific normalised difference | PSND | $(R_{800} - R_{470})/(R_{800} - R_{470})$ | (Blackburn, 1998b) |
| Plant water index | PWI | $R_{900}/R_{970}$ | (Peñuelas et al., 1994) |
| Renormalized difference vegetation index | RDVI | $(R_{800} - R_{670})/\sqrt{R_{800} + R_{670}}$ | (Roujean and Breon, 1995) |
| Zhen Index | Zen | $(R_{785} - R_{810})/((R_{785} + R_{810}) - (2 * R_{802}))$ | (Zhen et al., 2020) |
| Difference vegetation index | DVI | $R_{782}/R_{675}$ | (Peng et al., 2018) |
| Transformed soil adjusted vegetation index | TSAVI | $0.5 * (R_{782} - (0.5 * R_{675}) - 0.2)/((0.5 * R_{782}) + (0.5 * R_{675}) - 01)$ | (Baret et al., 1989) |
| Perpendicular vegetation index | PVI | $(R_{800} - (0.2 * R_{670}) - 06)/1.019)$ | (Darvishzadeh et al., 2006) |
| Ratio analysis of reflectance spectra chlorophyll c | RARSc | $(R_{760}/R_{500})$ | (Montesinos-López et al., 2017) |
| Green normalized difference vegetation index | GNDVI | $(R_{780} - R_{670})/(R_{780} + R_{670})$ | (Montesinos-López et al., 2017) |
| Red normalized difference vegetation index | RNDVI | $(R_{780} - R_{550})/(R_{780} + R_{550})$ | (Montesinos-López et al., 2017) |

Rxxx: Reflectance at wavelength 'xxx'.
Dxxx: First derivation of reflectance values at wavelength 'xxx'.
maxxxx: maximum value at wavelength 'xxx'.

Table S6: Selected single nucleotide polymorphisms based on BayesC using the posterior inclusion probability threshold of 0.10 for plant height (PH) under the B- management.

| Management | Trait | Chromosome | Marker ID | Minor allele frequency | Posterior inclusion probability |
|---|---|---|---|---|---|
| B- | PH | 1 | CM007647.1_6916155 | 0.38 | 0.814 |
| B- | PH | 1 | CM007647.1_115455586 | 0.12 | 0.264 |
| B- | PH | 2 | CM007648.1_14984692 | 0.44 | 0.277 |
| B- | PH | 2 | CM007648.1_43718939 | 0.12 | 0.162 |
| B- | PH | 2 | CM007648.1_108275267 | 0.47 | 0.112 |
| B- | PH | 2 | CM007648.1_112068617 | 0.49 | 0.128 |
| B- | PH | 2 | CM007648.1_112233172 | 0.49 | 0.291 |
| B- | PH | 3 | CM007649.1_57358057 | 0.31 | 0.733 |
| B- | PH | 4 | CM000780.4_162995251 | 0.29 | 0.387 |
| B- | PH | 4 | CM000780.4_174621437 | 0.33 | 0.215 |
| B- | PH | 4 | CM000780.4_214178316 | 0.36 | 0.133 |
| B- | PH | 4 | CM000780.4_238615869 | 0.26 | 0.677 |
| B- | PH | 5 | CM000781.4_4105099 | 0.32 | 0.143 |
| B- | PH | 5 | CM000781.4_17293217 | 0.28 | 0.137 |
| B- | PH | 6 | CM000782.4_93020758 | 0.08 | 0.122 |
| B- | PH | 6 | CM000782.4_168918815 | 0.33 | 0.190 |
| B- | PH | 7 | CM007650.1_161417373 | 0.30 | 0.187 |
| B- | PH | 8 | CM000784.4_67750993 | 0.12 | 0.172 |
| B- | PH | 9 | CM000785.4_28676437 | 0.23 | 0.217 |
| B- | PH | 9 | CM000785.4_28678507 | 0.22 | 0.211 |
| B- | PH | 9 | CM000785.4_32797796 | 0.49 | 0.146 |
| B- | PH | 9 | CM000785.4_32797855 | 0.48 | 0.210 |
| B- | PH | 9 | CM000785.4_150486056 | 0.30 | 0.176 |
| B- | PH | 10 | CM000786.4_133638870 | 0.40 | 0.139 |

Table S7: Selected single nucleotide polymorphisms based on BayesC using the posterior inclusion probability threshold of 0.10 for stalk diameter (SD) under the B- management.

| Management | Trait | Chromosome | Marker ID | Minor allele frequency | Posterior inclusion probability |
|---|---|---|---|---|---|
| B- | SD | 1 | CM007647.1_203994793 | 0.49 | 0.109 |
| B- | SD | 2 | CM007648.1_10595188 | 0.45 | 0.102 |
| B- | SD | 2 | CM007648.1_189507598 | 0.40 | 0.156 |
| B- | SD | 2 | CM007648.1_220144531 | 0.42 | 0.144 |
| B- | SD | 3 | CM007649.1_133566445 | 0.21 | 0.108 |
| B- | SD | 3 | CM007649.1_157011824 | 0.34 | 0.133 |
| B- | SD | 5 | CM000781.4_53086536 | 0.41 | 0.109 |
| B- | SD | 7 | CM007650.1_5874773 | 0.13 | 0.103 |
| B- | SD | 9 | CM000785.4_45942495 | 0.37 | 0.203 |

Table S8: Selected single nucleotide polymorphisms based on BayesC using the posterior inclusion probability threshold of 0.10 for plant height (PH) under the B+ management.

| Management | Trait | Chromosome | Marker ID | Minor allele frequency | Posterior inclusion probability |
|---|---|---|---|---|---|
| B+ | PH | 1 | CM007647.1_6540758 | 0.37 | 0.114 |
| B+ | PH | 1 | CM007647.1_200165533 | 0.37 | 0.210 |
| B+ | PH | 1 | CM007647.1_255269329 | 0.36 | 0.206 |
| B+ | PH | 2 | CM007648.1_129846748 | 0.40 | 0.265 |
| B+ | PH | 2 | CM007648.1_169473601 | 0.49 | 0.397 |
| B+ | PH | 3 | CM007649.1_158981956 | 0.22 | 0.109 |
| B+ | PH | 4 | CM000780.4_162995251 | 0.29 | 0.101 |
| B+ | PH | 4 | CM000780.4_176216697 | 0.15 | 0.114 |
| B+ | PH | 4 | CM000780.4_203497008 | 0.19 | 0.156 |
| B+ | PH | 5 | CM000781.4_3760644 | 0.45 | 0.412 |
| B+ | PH | 5 | CM000781.4_3760660 | 0.39 | 0.115 |
| B+ | PH | 5 | CM000781.4_75194167 | 0.18 | 0.137 |
| B+ | PH | 5 | CM000781.4_195847141 | 0.48 | 0.283 |
| B+ | PH | 6 | CM000782.4_108027757 | 0.49 | 0.121 |
| B+ | PH | 6 | CM000782.4_150545871 | 0.47 | 0.387 |
| B+ | PH | 6 | CM000782.4_168918815 | 0.33 | 0.239 |
| B+ | PH | 7 | CM007650.1_99946782 | 0.15 | 0.209 |
| B+ | PH | 8 | CM000784.4_155019631 | 0.23 | 0.228 |
| B+ | PH | 10 | CM000786.4_144153912 | 0.25 | 0.230 |
| B+ | PH | 10 | CM000786.4_145449487 | 0.20 | 0.144 |
| B+ | PH | 10 | CM000786.4_145449559 | 0.21 | 0.128 |

Table S9: Selected single nucleotide polymorphisms based on BayesC using the posterior inclusion probability of 0.10 for stalk diameter (SD) under the B+ management.

| Management | Trait | Chromosome | Marker ID | Minor allele frequency | Posterior inclusion probability |
|---|---|---|---|---|---|
| B+ | SD | 1 | CM007647.1_17324463 | 0.31 | 0.118 |
| B+ | SD | 1 | CM007647.1_49402080 | 0.27 | 0.120 |
| B+ | SD | 1 | CM007647.1_203994793 | 0.49 | 0.196 |
| B+ | SD | 1 | CM007647.1_286728045 | 0.47 | 0.149 |
| B+ | SD | 2 | CM007648.1_16421052 | 0.34 | 0.166 |
| B+ | SD | 2 | CM007648.1_32734863 | 0.26 | 0.143 |
| B+ | SD | 2 | CM007648.1_32734876 | 0.25 | 0.190 |
| B+ | SD | 2 | CM007648.1_38816460 | 0.26 | 0.306 |
| B+ | SD | 2 | CM007648.1_82458517 | 0.45 | 0.303 |
| B+ | SD | 2 | CM007648.1_169026437 | 0.36 | 0.447 |
| B+ | SD | 2 | CM007648.1_219293526 | 0.36 | 0.241 |
| B+ | SD | 3 | CM007649.1_205139465 | 0.29 | 0.113 |
| B+ | SD | 4 | CM000780.4_72636153 | 0.30 | 0.657 |
| B+ | SD | 4 | CM000780.4_164089694 | 0.10 | 0.119 |
| B+ | SD | 4 | CM000780.4_169305835 | 0.19 | 0.115 |
| B+ | SD | 4 | CM000780.4_193981325 | 0.29 | 0.112 |
| B+ | SD | 4 | CM000780.4_244379137 | 0.45 | 0.120 |
| B+ | SD | 6 | CM000782.4_2916367 | 0.16 | 0.114 |
| B+ | SD | 6 | CM000782.4_99963274 | 0.23 | 0.100 |
| B+ | SD | 6 | CM000782.4_103106825 | 0.12 | 0.118 |
| B+ | SD | 7 | CM007650.1_41779617 | 0.26 | 0.193 |
| B+ | SD | 7 | CM007650.1_41779668 | 0.37 | 0.563 |
| B+ | SD | 7 | CM007650.1_161417373 | 0.30 | 0.140 |
| B+ | SD | 8 | CM000784.4_151688268 | 0.29 | 0.102 |
| B+ | SD | 8 | CM000784.4_167863049 | 0.44 | 0.122 |
| B+ | SD | 9 | CM000785.4_18547071 | 0.20 | 0.115 |
| B+ | SD | 10 | CM000786.4_95462636 | 0.33 | 0.162 |

Table S10: Selected single nucleotide polymorphisms based on BayesC using the posterior inclusion probability threshold of 0.10 for shoot dry mass (SDM) under the B+ management.

| Management | Trait | Chromosome | Marker ID | Minor allele frequency | Posterior inclusion probability |
|---|---|---|---|---|---|
| B+ | SDM | 1 | CM007647.1_26874225 | 0.27 | 0.120 |
| B+ | SDM | 1 | CM007647.1_26874654 | 0.41 | 0.103 |
| B+ | SDM | 2 | CM007648.1_34931185 | 0.47 | 0.102 |
| B+ | SDM | 4 | CM000780.4_181569268 | 0.33 | 0.176 |
| B+ | SDM | 10 | CM000786.4_129809018 | 0.37 | 0.110 |

### Supplementary Figure

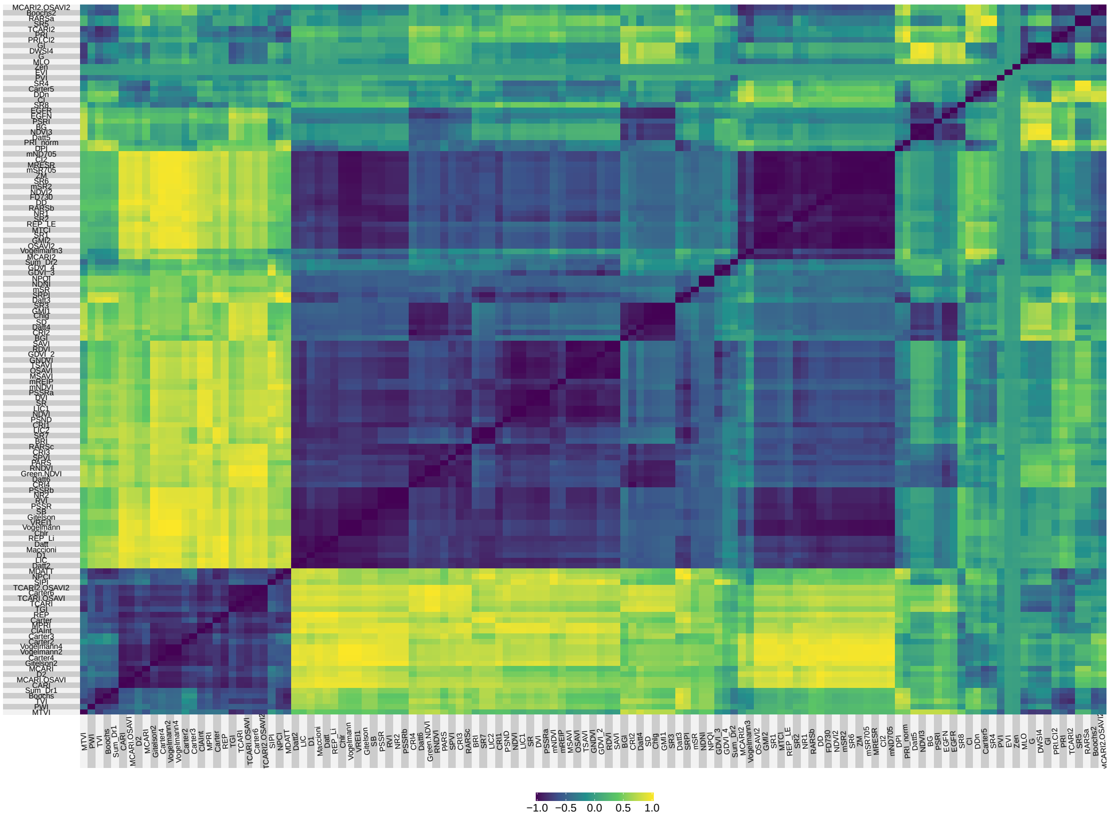


Figure S1: Correlation matrix of 131 hyperspectral indices. The correlation coefficients were computed from the averages of the two managements (B- and B+).
